# De novo haplotype reconstruction in viral quasispecies using paired-end read guided path finding

**DOI:** 10.1101/254987

**Authors:** Jiao Chen, Yingchao Zhao, Yanni Sun

## Abstract

**Motivation:** RNA virus populations contain closely related but different viral strains infecting an individual host. As the selection acts on clouds of mutants rather than single sequences, these viruses have abilities to escape host immune responses or develop drug resistance. Reconstruction of the viral haplotypes is a fundamental step to characterize the virus population, predict their viral phenotypes, and finally provide important information for clinical treatment and prevention. Advances of the next-generation sequencing technologies open up new opportunities to assemble full-length haplotypes. However, error-prone short reads, high similarity between related strains, unknown number of haplotypes pose computational challenges for reference-free haplotype reconstruction. There is still big room to improve the performance of existing haplotype assembly tools.

**Results:** In this work, we developed a *de novo* haplotype reconstruction tool PEHaplo for viral quasispecies data, which contains a group of related but different viral strains. PEHaplo employs paired-end reads to distinguish highly similar strains. We applied it to both simulated and real quasispecies data, and the results were benchmarked against several recently published haplotype reconstruction tools. The comparison shows that PEHaplo outperforms the benchmarked tools in a comprehensive set of metrics.

**Availability:** The source code and the documentation of PEHaplo is available at https://github.com/chjiao/PEHaplo.

## 1 Introduction

High mutation rate, nature selections, and recombination can lead to high genetic diversity of RNA virus populations (Domingo-Calap *et al*., 2016), which consist of closely related but different viral strains. These groups of virus populations are often termed as viral quasispecies (Nowak, 2006). Each strain in quasispecies is defined by its haplotype sequence. Commonly known examples of the fast mutating viruses include clinically important viruses such as human immunodeficiency virus (HIV-1) and the hepatitis C virus (HCV). The genetic heterogeneity of the virus populations is key to their adaptive behavior. As the selection works on a set of sequences rather than one, high genetic diversity gives the viruses the abilities to escape host immune responses or develop drug resistance. Reconstruction of the viral haplotypes is a fundamental step to characterize the structure of the virus populations, predict viral phenotypes, and finally provide important information for clinical treatment and prevention (Schirmer *et al*., 2012).

Development of next-generation sequencing technologies sheds light on characterizing the haplotypes and their abundance in heterogeneous virus populations. The deep sequencing of virus population samples becomes available and various methods and tools have been developed for viral haplotype reconstruction (Baaijens *et al*., 2017). The methods can be divided into two groups based on their dependency on a reference genome (Beerenwinkel *et al*., 2012). The first group of methods need reference genomes and employ read alignments against the reference sequence to infer haplotypes. However, due to the high mutation rate, high quality reference genomes of a virus population are not always available. In particular, for emerging infectious viral diseases such as SARS that lack reference genomes during the breakout, reference-based methods are not plausible. The second group of methods belong to *de novo* haplotype reconstruction, which do not require reference genomes. This type of method can be applied to characterize new viral strains or novel haplotypes. Our work belongs to the second group.

A recent review of chosen haplotype reconstruction tools has shown that haplotype recovery is a computationally challenging problem (Schirmer *et al*., 2012). The authors’ benchmarking results on a series of data demonstrated that the performance of the tested programs is poor when sequence divergence is low. In addition, these programs failed to recover rare haplotypes. Thus, there is a need for new methods and tools for more accurate haplotype reconstruction.

The tested programs in the review (Schirmer *et al*., 2012) all belong to group 1, which need alignments against a reference sequence as input. Without using a reference sequence, our method of haplotype reconstruction from deep sequencing data takes a method similar to *de novo* genome assembly. Applying assembly to viral haplotype reconstruction faces several challenges. The first challenge is to distinguish highly similar genomes of different strains. Figure S2 in the Supplementary Materials presents a partial multiple alignment of five haplotypes in a mixed HIV sample, showing the high sequence similarity between the genomes of interest. Existing assembly methods tend to produce either short or chimeric contigs for deep sequencing data containing highly similar genomes. Second, it is difficult to distinguish sequencing errors from mutations of a rare haplotype. Third, reads originating from haplotypes of low abundance tend to have small overlaps and thus only fragments of haplotypes can be reconstructed.

### 1.1 Related work

There exist a number of metagenomic assembly tools that could be applied to assemble viral genomes in quasispecies data (Namiki *et al*., 2012; Treangen *et al*., 2013; Laserson *et al*., 2011; Peng *et al*., 2011; Luo *et al*., 2012; Salzberg *et al*., 2008; Wu *et al*., 2012). However, these assembly tools are not designed to distinguish different haplotypes and only produce fragmented or chimeric contigs.

There are a group of tools designed specifically for haplotype reconstruction (Zagordi *et al*., 2011; Malhotra *et al*., 2013; Prabhakaran *et al*., 2010; Töpfer *et al*., 2013; Prosperi and Salemi, 2012; Astrovskaya *et al*., 2011; T O’Neil and Emrich, 2012; Huang *et al*., 2011). Of them, HaploClique (Töpfer *et al*., 2014), SAVAGE (Baaijens *et al*., 2017), and MLEHaplo (Malhotra *et al*., 2015) were recently published and are highly related to ours. Like our work, all of them utilized paired-end reads. HaploClique uses the insert size distribution for detection of large indels. However, it needs a reference sequence for generating alignment graphs. HaploClique provides a source of inspiration for SAVAGE, which is the first tool for *de novo* assembly of viral haplotypes using overlap graphs. SAVAGE also took advantage of paired-end reads and merge short reads using cliques. The authors benchmarked SAVAGE with other virus assembly tools and showed that SAVAGE outperformed other tools in a comprehensive set of assembly metrics. MLEHaplo also explicitly employs paired-end reads for finding top-score paths. MLEHaplo and our method are based on two different graph models: de Bruijn graph and overlap graph. Thus, different error correction and graph pruning techniques are applied. In addition, during path finding, we carefully distinguish paired-end connections formed by different types of nodes in order to improve the accuracy of the chosen paths. As we focus on *de novo* assembly tools, we will benchmark our work against SAVAGE and MLEHaplo.

In this work, we designed and implemented PEHaplo, which assembles viral haplotypes from deep sequencing data. Sequence assembly has been an intensive research area and new methods or implementations are emerging quickly. We created a novel overlap graph incorporating paired-end reads information. The paired-end information is utilized in both graph pruning, path finding, and contig refinement. We applied PEHaplo to both simulated and real viral deep sequencing data and compared the assembly performance with the recently published tools. The experimental results show that PEHaplo can recover viral haplotypes with longer contigs and higher accuracy.

## 2 Methods

A major challenge for viral haplotype reconstruction is the high sequence similarity between viral strains. In particular, the distribution of the mutations/insertions/deletions largely determines the difficulty levels of the problem. Here, we use LCS (longest common substring) to refer to the longest common substring between any two neighboring mutations/insertions/deletions. Depending on the size of the LCS, we have three cases as shown in Figure 1.

- If LCS size ≤ read size, haplotype reconstruction can be solved based on read overlaps (Figure 1 (A)).
- If LCS size ≥ read size but ≤ insert size, paired-end reads are able to distinguish different haplotypes (Figure 1 (B)).
- If LCS size ≥ insert size, coverage information can be utilized to distinguish haplotypes of different abundance (Figure 1 (C)).

In order to classify viral haplotype reconstruction problems in the above three cases, it is ideal to know the LCS distributions inside each quasispecies. While the insert size can be estimated for different sequencing platforms, it is not feasible to empirically obtain all viral strains and compute the sizes of their LCSs. We thus rely on quasispecies theory (Nowak, 2006) for estimating the LCS sizes using an average viral mutation rate. The detailed method and also the generated distribution of LCSs can be found in Supplementary Materials Section 1.

According to our computed LCS distribution, the LCS sizes span all three cases in Figure 1. Thus, our methods use three types of information for virus assembly. 1) Paired-end reads. As paired-end reads are sequenced from the same fragment, they thus belong to the same haplotype. 2): Coverage. If two strains have highly different coverages, they can be distinguished using coverage information. 3): Enumeration of cliques. Reads forming cliques in the overlap graph tend to come from the same haplotype and thus can be merged as a super-read. This process can be iteratively applied to extend local haplotype to global one and is an important component of several recently published tools (Töpfer *et al*., 2014; Malhotra *et al*., 2015; Baaijens *et al*., 2017). Although paired-end reads have been employed in haplotype reconstruction previously, they were not carefully elaborated and analyzed. We conducted a deep analysis of the utility and limitations of using paired-end reads for haplotype reconstruction.

The remainder of the Methods Section is organized as follows. We will first introduce the paired-end overlap graph and the key idea of path finding using paired-end reads. Then we will show the complete pipeline and describe the major components.

**Fig. 1.**
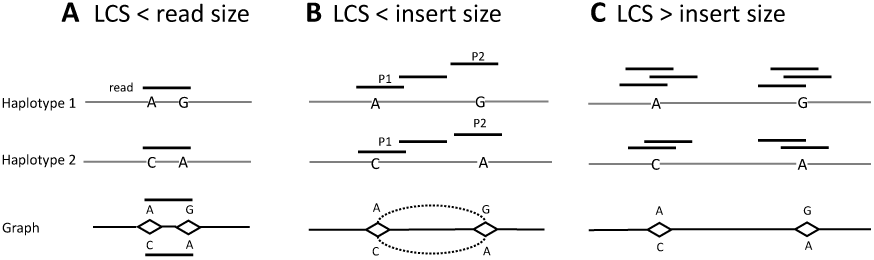
Distinguishing two haplotypes with different LCS (longest common substring) sizes. In panel B, P1 and P2 represent the ends of a read pair. The problem becomes harder with increase of the LCS size.

### 2.1 Paired-end overlap graph and path finding

An overlap graph *G*(*V, E*) is a weighted directed graph that reflects overlaps between reads. Each node *v* ∈ *V* represents a read. An overlap between two reads is formed if the suffix of a read matches the prefix of another read. Given any two reads *r*_1_, *r*_2_, and an overlap threshold *l*, if the overlap size between *r*_1_ and *r*_2_ is greater than *l*, a directed edge is added from the nodes representing *r*_1_ and *r*_2_ in *G*. The edge weight is the overlap size.

In our method, we constructed a paired-end overlap graph (PE_G), which adds information from paired end reads to a standard overlap graph. PE_G has the same node set as an overlap graph but it has two sets of edges. One set of edges are inherited from a standard overlap graph. The other set of edges connect nodes whose reads form paired-end reads. Intuitively, while an overlap graph records the connectivity between reads, PE_G also records the number of paired-end reads between nodes. Thus, PE_G can be defined as *G*(*V, E, E’*), where *E* is the same as in an overlap graph. If two reads form a paired-end read pair, an edge in set *E’* is created between the corresponding nodes.

Figure 2 shows an example of paired-end overlap graph. Figure 2(A) contains two strains with only two mutations and also the reads sequenced from them. The overlap threshold is set as half of the read size. The paired-end overlap graph PE_G is constructed using the reads and is shown in (B). The edges in *E* are shown using solid lines while the edges in *E’* are shown using dashed lines. Nodes a.1 and a.2 are a read pair and thus form an edge in *E’*. Similarly, nodes d.1 and d.2 have an edge in *E’* because d.1 and d.2 are a read pair.

In the graph, there are four complete paths: *a*.1→ *b* → *c* → *e* → *f* → *a*.2, *a*.1→ *b* → *c* → *e* → *f* → *d*.2, *d*.1 → *b* → *c* → *e* → *f* → *a*.2, and *d*.1→ *b* → *c* → *e* → *f* → *a*.2. The goal of assembly is to output the two correct paths (i.e. *a*. 1 to *a*.2 and *d*.1 to *d*.2). The edges in *E’* will guide the extension of a path that is composed of edges in *E*. Specifically, for a path starting with a.1, it will decide whether to extend to a.2 or d.2 at node f. The dashed edges in *E’* (*a*.1 → *a*.2) will guide the path to the correct node a.2. Similarly, a path starting with d.1 will end with d.2 based on the guidance of the dashed edge *d*.1 → *d*.2. Thus, the path finding using the paired-end information will output two paths, representing the two haplotypes.

**Fig. 2.**
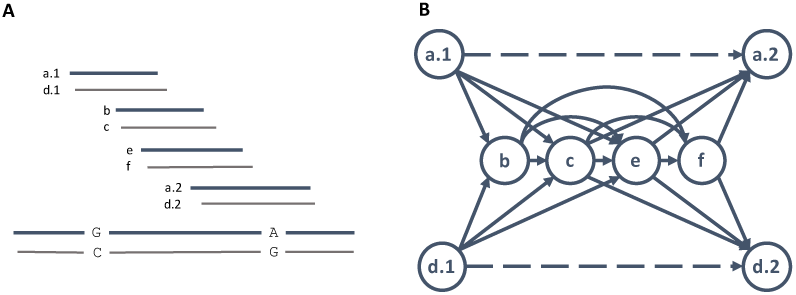
(A) The bottom two long lines represent two haplotypes, which only differ by two mutations at two loci (G-C and A-G). Short lines represent reads sequenced from the two strains. The reads are sorted by their read mapping positions against their native strain, a.l and a.2 are a read pair from the thick strain, d.l and d.2 are the read pair from the thin strain. (B) Paired-end overlap graph. Nodes b, c, e, and f originate from the common region of the two strains. The dashed lines represent paired-end read connection.

### 2.2 The whole pipeline of PEHaplo

There are five major components in the pipeline of PEHaplo as shown in Figure 3. In the first pre-processing stage, reads with low-quality or ambiguous base calls are filtered or trimmed. Base-calling errors or indels are corrected from the filtered set of reads using alignment-based error correction. Duplicated reads and substring reads are then removed from the corrected reads. The detailed pre-processing description can be found in the Supplementary Materials Section 2. Second, an overlap graph is built from the pre-processed reads and the strand of reads are adjusted by traversing the graph. The detailed strategy about strand adjustment can be found in Supplementary Materials Section 3. The output reads will have the same orientation. The third stage will build the overlap graph again from the strand-adjusted reads and utilize various graph pruning methods to remove possible random overlaps and simplify the graph for efficient assembly. In the fourth stage, *E’* will be constructed and paired-end guided path finding algorithms are applied to produce contigs from the paired-end overlap graph. Finally, we align paired-end reads against produced contigs to identify and correct potential mis-join errors.

**Fig. 3.**
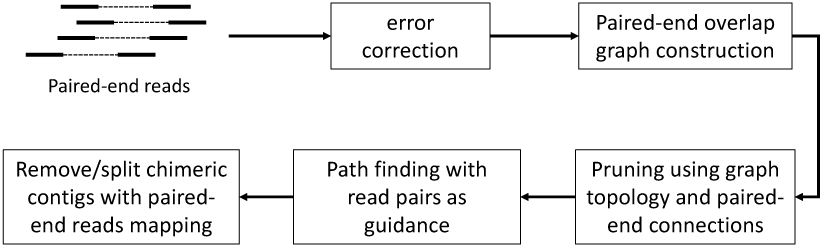
The pipeline and main components of PEHaplo. Note that the error correction component is implemented using Karect (Allam *et al*., 2015), which uses alignments between reads rather than alignments between reads and a reference genome for error correction.

#### 2.2.1 Paired-end overlap graph construction

There are two steps in construction of the paired-end overlap graph. In the first step, we construct the standard overlap graph, collapse nodes, and merge cliques. In the second step, we identify the number of paired-end reads between nodes in the overlap graph and remove false connected edges using added paired-end information. This section will focus on the overlap computation between reads.

All reads remained after pre-processing are used to construct the overlap graph. A straightforward overlap detection method requires *O*(*n*^2^) comparisons, which is computationally expensive for large sequencing data sets. There are efficient implementation of all-pairs suffix-prefix comparison algorithms based on data structures such as hashing table or compact prefix tree (Gonnella and Kurtz, 2012; Haj Rachid and Malluhi, 2015). In PEHaplo pipeline, we first compute all suffix-prefix matches between reads for read orientation adjustment, and then compute again for the final overlap graph construction.

**Overlap cutoff estimation** The overlap cutoff *l* is an important parameter. A small *l* tends to keep most true overlaps but also introduces more false connected edges, while large *l* is likely to eliminate most false overlaps but can possibly miss true connections for reads from lowly sequenced regions. We use exponential distribution to estimate the appropriate overlap cutoff.

Let *N* be the total reads number, *r* be the read length and *L* be the genome size, the sequencing coverage is thus calculated as *C* = *Nr*/*L*. We use the Poisson distribution to model the number of reads sequenced from unit length of a genome. The parameter λ in the Poisson distribution is estimated as *N/L* (Wang *et al*., 2009). The distance (*X*) (see Figure 4) between two adjacent reads will thus follow an exponential distribution. The corresponding cumulative distribution function is the probability that two adjacent reads have an overlap size of at least *r* – *d*. For example, Let *r* = 250, *L* = 10, 000, and *N* = 800. Then the sequencing coverage *C* is 20 and *λ*(*N/L*) is 0.08. Thus, for given d as 70, we have *F*(*X* ≤ 70) = 0.9963. That is, there is 99.63&&#x0023;x0025; possibility that two adjacent reads will form an overlap with size of at least 180 (i.e. 250-70). Since we have 800 reads, the expected number of pairs with overlap smaller than 180 is: 799 * (1 — 0.9963) = 2.96. Therefore, we may have four contigs for the genome under this overlap cutoff. While keeping the connectivety of the graph, the overlap threshold should be as large as possible.

**Fig. 4.**
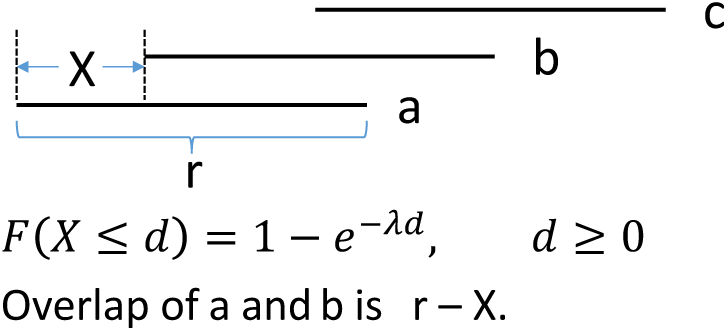
The distance between two adjacent reads can be estimated by an exponential distribution. *a,b, c* represent three reads, *r* is the read length.

#### 2.2.2 Graph pruning

The original overlap graph is usually very complex because of the large data size, transitive edges, sequencing errors, and highly similar regions shared by haplotypes. Besides commonly used transitive reduction and node collapsing (Yuan *et al*., 2015), we apply an iterative graph pruning procedure to repeatedly simplify the graph at each iteration.

**Merge reads in cliques** We are interested in cliques in the overlap graph because reads within a clique can share true mutations while sequencing errors are usually more random and are not shared by the majority of reads. Therefore, cliques can be used to distinguish true mutations from sequencing errors. Several recent haplotype reconstruction tools (Baaijens *et al*., 2017; Töpfer *et al*., 2014) merge reads inside cliques as super-reads and conduct iteratively haplotype extension. In our methods, we apply the step of merging nodes inside of cliques.

**Removing false edges using read pairs** Due to the nature of viral quasispecies, different haplotypes of one species usually have very high sequence similarity (possibly over 90%), which can easily cause overlaps between reads originating from different strains. Therefore, having a suffix-prefix match does not guarantee that the two reads originate from the same viral strain. Wrong edges increase the complexity of the graph and may also produce misjoined contigs. We employ paired-end information to remove potentially wrong edges.

The key idea is that for an edge formed by reads from different haplotypes, its two end nodes usually have other incident edges incurred by the correct connections. The wrong edge or the contigs containing the wrong edge are not well supported by read pairs. Therefore, we use paired-end information as evidence to remove false edges. We examine each edge formed by nodes with large in or out degrees (Cormen, 2009) because it is likely that some of the incident edges are false overlaps.

Figure 5(A) presents a case. Edge *u* → *v* is one of the many edges incident to nodes *u* or *v*. We apply the following rules for edge *u* → *v*: if there is no read pair support between *u* and *v*, between *u’s* predecessor nodes and *v*, or between *u* and *v’s* successor nodes, and the sequence formed by joining *u, v* is longer than an insert size cutoff, we will remove *u* → *v*. The insert size cutoff can be customized depending on the given data properties. To remove false connected edges, we need to traverse each node and edge of the overlap graph. The time complexity is *O*(*V* + *E*).

#### 2.2.3 Paired-end guided path finding

Once the overlap graph is pruned, paired-end connections will be added and thus form the paired-end overlap graph PE_G. As aquickreview, PE_G is defined as (*V, E, E’*), where *E* contains the edges from read overlaps while *E’* contains edges from paired-end connections. The weight of an edge in *E’* represents the number of paired-end read pairs between two end nodes. Note that after node collapse, each node can contain multiple reads.

The problem of assembling a single haplotype in the graph PE_G can be formulated as finding a path *p*, where each edge in *p* ∈ *E*, so that *p’s* weight defined using edges in *E’* is maximized. Intuitively, we look for paths with the best support of paired-end connections. This process can be repeated to find *k* longest paths, whose path weight is defined by edge weight in *E’*. However, we can prove that finding the path with the most number of paired-end connections in PE_G is NPC. The detailed proof can be found in the Supplementary Materials Section 4.

**Fig. 5.**
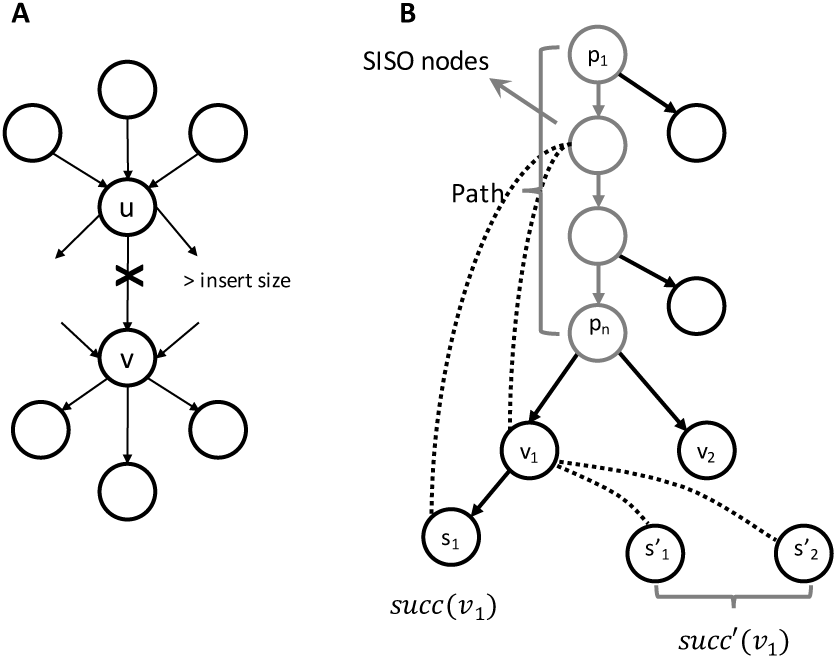
(A) Removing false edges using paired-end information. The overlap edges with cross will be removed if insufficient read pairs exist between their ends. (B) In this example, the ending node *p_n_* of current path has two successors *v*_1_ and *v*_2_. The path nodes and a SISO node in the path are marked. The solid lines are overlaps and dashed lines are pairedend connections between nodes. Six scores are computed to select the right successor for extending the path. Among which, five are calculated as paired-end edge weights between nodes in current path and nodes associated with the successor. As for *v*_1_, *succ*(*v*_1_) and *succ’*(*v*_1_) are shown in the figure. The group 1 scores contain the paired-end edge weights between the SISO node and *v*_1_, *s*_1_, and 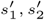. Group2 scores contain the paired-end edge weights between path nodes {*p*_1_,…, *p_n_* – 1} and *v*_1_, *s*_1_.

In our method, we use a DFS-based path extension algorithm for path finding and use a greedy algorithm to select the right node for path extension. At each vertex with multiple successors, we apply the greedy algorithm to choose the best node for extension. The greedy algorithm carefully considers paired-end connections between different types of nodes and also the coverage information. Paired-end connections between nodes can be efficiently accessed from the constructed paired-end graph PE_G.

**Scores calculation for path extension** To find correct paths from the graph, the right node need to be selected each time we extend the path. In particular, when a node has multiple successors, a right choice must be made for path extension. In general, we choose a successor with the most paired-end connections to the current path. Different types of paired-end connections were treated with different priorities in distinguishing haplotypes. In particular, single-in single-out (SISO) nodes are differentiated from other nodes. SISO nodes have in-degree of 1 and out-degree of 1 and thus are unique to one haplotype. Any paired-end connection incident to SISO nodes can be used to recruit nodes that belong to the same haplotype. On the contrary, nodes originating from common regions of two or more haplotypes are not SISO nodes. Paired-end connections to those nodes do not provide useful guidance in path finding. In our algorithm, we distinguish paired-end connections involving SISO nodes and other nodes.

**Algorithm 1.**
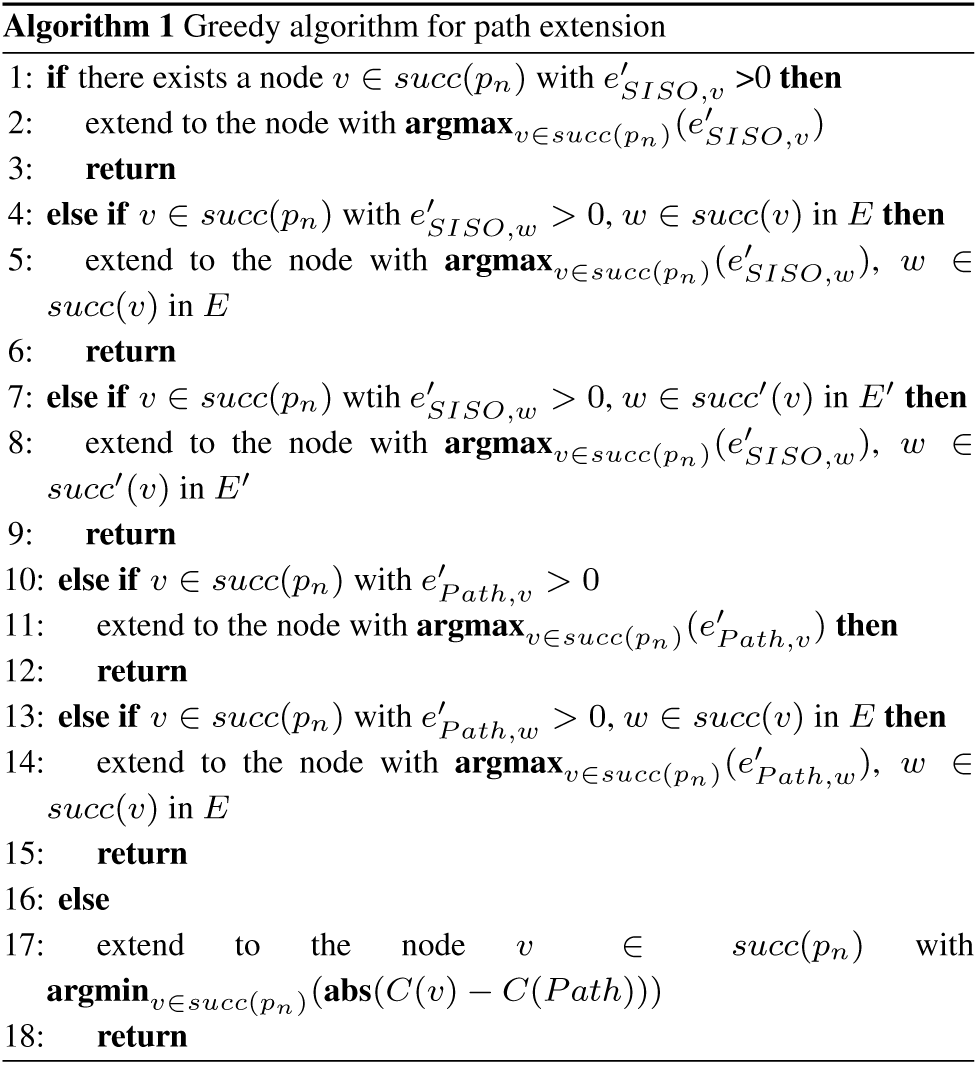
Greedy algorithm for path extension.

For *e’* ∈ *E’*, *e’*(*u, v*) is the edge weight between *u* and *v* in *E’*. Let *Path* = {*p*_1_,*p*_2_,…,*P_n_*} be the current path. The ending node *p_n_* in the current path has multiple successors. As we have two sets of edges in PE_G, let *succ*(*v*) in *E* represents *v’s* successor nodes in the standard overlap graph, *succ*(*v*) in *E’* represents all nodes that form paired-end connections with *v*. For each successor *v* of *p_n_* (*v* ∈ *succ*(*p_n_*)), we compute six scores, which are divided into three groups. An example of the scores calculation is shown in Figure 5(B). Group 1 contains the paired-end edge weights between SISO nodes in *Path* and *v* (score 1), *v’s* successor nodes in *E* (score 2), and *v’s* successor nodes in *E’* (i.e. all nodes that form paired-end connections witht), score 3). Group 2 contains the paired-end edge weights between all path nodes (except *p_n_*) and *v* (score 4), *v’s* successor nodes in *E* (score 5). The third group contains coverage difference between *Path* and *v* (score 6).

The pseudocode in **Algorithm 1** describes the greedy algorithm that chooses the locally optimal node for path extension based on the above scores. 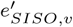 represents a paired-end edge weight between SISO node(s) in the current path and *v*, which is a successor node of *p_n_*. Function *C*(*v*) denotes the read coverage of node *v*.

#### 2.2.4 Correcting contigs with paired-end read distribution

To further improve the quality of assembled contigs, we apply a contig correction method similar to the tool PECC (Li *et al*., 2017). With the contigs generated after path finding, we align the raw reads to them and split contigs from the locations with low read pairs coverage (Figure 6).

**Fig. 6.**
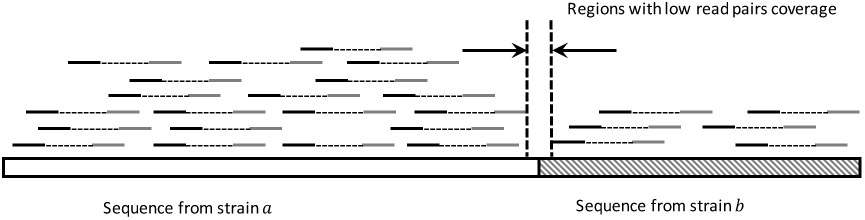
Read pairs mapping profile on a misjoined contig. The contig is shown as the long bar at the bottom, which is misjoined with two sequences from strains *a* and *b*. The dashed line connects the two ends of a read pair. Fewer read pairs will go across the misjoined location, thus revealing a valley in the aligned reads profile, which can be used to split the contig.

## 3 Results

We have designed and implemented PEHaplo, a *de novo* viral quasispecies assembly method that uses paired-end reads to guide path finding in paired-end overlap graph. In this section, we will evaluate the performance of our method on both simulated and real viral quasispecies data sets. The simulated data set includes both the commonly adopted HIV data set and also highly biased data set with rare haplotypes. For each experiment, we will present the performance of PEHaplo and benchmark it with recently published quasispecies *de novo* assembly tools. In addition, we carefully evaluate the performance of each main component in the whole pipeline. The results show that our tool produces fewer and significantly longer contigs that can recover a majority of the haplotypes.

### 3.1 Experimental data sets

We evaluated PEHaplo on several simulated data sets, one real HIV-1 Illumina MiSeq sequencing data set, and one real Influenza sample. Both the simulated and real HIV-1 data sets were generated from a mixture of five well-studied HIV-1 strains (HXB2, JRCSF, 89.6, NL 43 and YU2). These strains have pairwise sequence similarities from 91.8% to 97.4% (Supplementary Table S1). HXB2 and NL43 have the highest similarity with LCS of size 427bp (Supplementary Table S2). We choose HIV because it is adopted by other viral haplotype reconstruction tools and has become a gold standard for performance evaluation.

### 3.2 Evaluation metrics

As the haplotype sequences and compositions are known in these datasets, we are able to evaluate the quality of the assembled contigs generated by all tools. We compared the produced results to recently published *de novo* assembly tools IVA (Hunt *et al*., 2015), MLEHaplo (Malhotra *et al*., 2015) and SAVAGE (Baaijens *et al*., 2017).

Following SAVAGE, we use a third-party tool MetaQuast (Mikheenko *et al*., 2015) for evaluating the output of all tested tools. MetaQuast integrated several components for convenient *de novo* assembly performance evaluation. It aligns the generated contigs to the viral reference genomes and reports the number of contigs, N50, unaligned length, target genome(s) covered, mismatch and indel rates, etc. N50 length is defined as the maximal length that all contigs of at least this length contain at least 50% of all the contig bases. A contig can be partially aligned to a reference sequence. Thus, the total length of all unaligned parts are reported as “unaligned length”. For the aligned parts, “target genome(s) covered” and “mismatch and indel rates” are computed. Target genome(s) covered is the percentage of reference genomes that are aligned by contigs, and mismatch/indel rate is the percentage of mismatchs/indels of aligned contigs.

### 3.3 Results on HIV simulated data set

We first applied PEHaplo on a simulated HIV-1 quasispecies data set. We used ART-illumina (Huang *et al*., 2012) to simulate 1.9e+5 paired-end, 250bp error-containing MiSeq reads from the five HIV-1 strains with average insert size of 600 bp and standard deviation of 150 bp. The total coverage of the 5 strains is ∼5000x, which is close to the coverage of real HIV quasispecies data commonly used by existing tools. To obtain a more realistic data set, a fitness based power law equation (Barbosa *et al*., 2012) was used to simulate the coverage distribution among five strains: 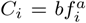, where *C_i_* and *f_i_* denote the coverage and fitness of strain *i*, respectively. The coverages for each strain in the simulated data set are: 89.6 - 2190x, HXB2 - 1095x, JRCSF - 730x, NL43 - 547x, YU2 - 438x.

Following the PEHaplo pipeline, we first performed error correction and duplicated sequence removal on the raw simulated data set. With 1.9e+5 error corrected reads, 48,833 reads were kept after removing duplicates. Only those reads that duplicate at least 3 times in the raw data were kept, further reducing the reads number to 26,961. After adjusting reads orientation, an overlap graph was constructed with the tool Apsp (Haj Rachid and Malluhi, 2015). The original overlap graph has 26,961 nodes and 977,570 edges. After merging cliques, removing transitive edges and collapsing nodes, 63 nodes and 67 edges were left. We then applied false edge removal and node collapsing on the graph, and further reduced it to 48 nodes and 44 edges. Paths and contigs were generated from this pruned graph using the greedy algorithm described in the Methods Section.

To evaluate the effectiveness of our false edges removal step, we also assembled the simulated reads without the false edges removal step. The results are shown in Supplementary Table S3. Comparing to contigs with false edges removal, the genome covered fraction is reduced from 97% to 91.8%. Also, the N50 value is decreased while the mismatch rate is increased. The results reveal that our false edges removal step is effective and can improve the final results.

PEHaplo generated 12 contigs from the simulated data set and the results are summarized in Table 1. The contigs are able to cover over 96% on the 5 viral strains, with aN50 of 9,262 bp. The largest contig has a length of 9,668bp, which can almost cover a complete HIV strain. Meanwhile, these contigs have low mismatch and indel rates.

We also assembled the simulated reads with benchmark tools IVA, MLEHaplo, and SAVAGE and summarized their results in Table 1. With the default parameters, IVA produced a single, long contig from the error corrected reads. This long contig has a length of 13,434 bp and can cover the whole genome of the strain 89.6 and about 43% of the strain HXB2. The results of IVA reveal that it tends to generate one consensus genome sequence corresponding to the haplotype with the highest coverage. Other strains are largely missed. Using k-mer size of 55, MLEHaplo produced 205 contigs that cover 78% of the five HIV-1 strains. The contigs it produced are quite fragmented, with a low N50 value of 671 bp and the longest contig of 1,716 bp. Following the guidance of SAVAGE tutorial, we set the overlap cutoff as 180 bp. SAVAGE produced 64 contigs that cover over 97% of the reference genomes, with a N50 of 1,926 bp, and the largest contigs of 7,941 bp.

Comparing to IVA and MLEHaplo, PEHaplo is able to produce longer contigs with fewer mismatches and indels on the simulated HIV-1 data set. SAVAGE also shows better performance than IVA and MLEHaplo on this data set. But it produced many short contigs (N50 value 1,926 bp vs. 9,262 bp of PEHaplo). Some of them cannot be aligned to the reference genomes, leading to larger “unaligned length”.

#### 3.3.1 Paired-end reads guided path finding is able to generate accurate long contigs

The greedy algorithm that carefully utilizes paired-end information plays a crucial role for producing high quality long contigs. In this section, we focus on evaluating the performance of path finding and investigating whether the improved performance of PEHaplo is simply due to the pruned graph or the combination of the pruned graph and path finding algorithm. Thus we applied popular *de novo* metagenomic assembly tools IDBA-UD (Peng *et al*., 2012) and Ray Meta (Boisvert *et al*., 2012) on the pruned overlap graph that used as the input for path finding in PEHaplo. In addition, as SAVAGE is the newest viral haplotype reconstruction tool and has better performance than IVA and MLEHaplo, we also applied SAVAGE on the same pruned overlap graph as PEHaplo.

With the same input reads or graph, we compared the output of these tools using MetaQuast and presented the results in Supplementary Table S3. The results revealed that those contigs assembled by IDBA-UD, Ray Meta and SAVAGE from the reduced overlap graph are fragmented and could only cover a small proportion of the five reference genomes. These contigs have low rate of mismatches and indels, but their average lengths are much shorter than PEHaplo. The experiments show that the paired-end guided path finding algorithm in PEHaplo is essential for producing long haplotype segments from the viral quasispecies sequencing data.

**Table 1.** Assembly results on simulated HIV data set for IVA, MLEHaplo, SAVAGE and PEHaplo. Contigs that are at least 500 bp are aligned to the reference haplotype sequences with a similarity cutoff of 98%.

| Tools | Contigs num | N50 | Genomes covered (%) | Unaligned length (bp) | Mismatch rate(%) | Indels (%) |
| --- | --- | --- | --- | --- | --- | --- |
| IVA | 1 | 13,434 | 28.7 | 0 | 0.809 | 0.051 |
| MLEHaplo | 205 | 671 | 78.0 | 81,125 | 0.542 | 0.008 |
| SAVAGE | 64 | 1,926 | **97.32** | 4792 | **0.009** | 0.004 |
| PEHaplo | 10 | 9,274 | 97.0 | **0** | 0.026 | **0.002** |

### 3.4 Benchmark on HIV MiSeq data set

To further assess the performance of assembly methods, we applied PEHaplo on a real HIV quasispecies data set (SRR961514) sequenced from the mix of five HIV-1 strains with Illumin MiSeq sequencing technology (Di Giallonardo *et al*., 2014). This data set contains 714,994 pairs (2×250 bp) of reads that cover the five strains to 20,000x.

We used PEHaplo to perform similar pre-processing procedures on the real HIV quasispecies data. With 774,044 filtered and error corrected reads, 98,947 reads were kept after removing duplicates and substrings. Since the raw data set has extremely high coverage on the five strains, we still kept those reads that duplicate at least three times in the raw data set. After these pre-processing procedures, 26,691 reads were kept for strand adjustment and assembly.

PEHaplo produced 33 contigs from the real MiSeq HIV data set that can cover over 92% of the five HIV-1 strains. These contigs have a N50 value about 2,500 bp and the longest contig is 9,108 bp. The results are summarized in Table 2. Compared to simulated HIV data set, PEHaplo has generated more contigs but with a lower N50 value and higher mismatches and indels on the real data set. We notice that the real HIV data set contains more sequencing errors and has a larger variation for insert size than the simulated data set.

We again compared the performance of PEHaplo with IVA, MLEHaplo and SAVAGE. IVA generated 10 contigs that can cover about 20% of the five strains. Similar to the simulated data set, these contigs still cover larger parts on haplotypes with higher sequencing coverage. With the same parameters as before, MLEHaplo produced 234 contigs that can cover over 53% of the five genomes with similar mismatch and indel rates to the simulated data set. It generated much longer contigs on the real data, with a N50 value of 6,501 bp and the largest contig of 8,470 bp. However, these contigs contain many misjoined segments. Over 150 contigs with total length of 787,272 cannot align to any reference genomes. Since the SAVAGE paper (Baaijens *et al*., 2017) has shown their results on the same data set, we use the metrics in their literature for evaluation. From their results, SAVAGE produced 846 contigs that cover over 92% of the reference genomes, with a N50 of 588 bp, and largest contig of 1,221 bp (Table 2).

On the real HIV data set, PEHaplo can still produce longer contigs with fewer mismatches than all three benchmarked tools. Overall, PEHaplo is able to assemble short reads that are sequenced from multiple viral strains sharing high similarities, generating long, high quality contigs that can reconstruct most of the target haplotypes. In Supplementary Figure S2, we show the contig alignment result on HXB2 strain for PEHaplo and SAVAGE. This figure clearly shows that our tool usually produces fewer but longer contigs.

**Table 2.** Assembly results on real HIV MiSeq data set for IVA, MLEHaplo, SAVAGE and PEHaplo.

| Tools | Contigs num | N50 | Genomes covered (%) | Unaligned length (bp) | Mismatch rate (%) | Indels (%) |
| --- | --- | --- | --- | --- | --- | --- |
| IVA | 10 | 1,150 | 20.1 | 1150 | 0.660 | 0.052 |
| MLEHaplo | 234 | 6,501 | 53.6 | 786,272 | 0.588 | **0.035** |
| SAVAGE | 846 | 588 | **92.6** | **0** | 0.161 | 0.040 |
| PEHaplo | 24 | 2,223 | 92.98 | **0** | **0.016** | 0.045 |

### 3.5 Benchmark on simulated biased HIV data sets

It is a major challenge to reconstruct the low-abundance haplotypes in quasispecies data. To evaluate the performance of our methods on assembling low abundance strains, we used HIV strains HXB2 and NL43 to simulate three groups of data sets with extremely biased coverages. We choose HXB2 and NL43 because they share the highest similarity and longest common region among five HIV strains, representing the hardest case for assembly. The total coverage for each group is 1000x, with HXB2-900x, NL43-100x; HXB2-950x, NL43-50x; and HXB2-990x, NL43-10x for each group, respectively. These data sets contain 250 bp paired-end reads produced by ART-illumina with average insert size of 600 bp and standard deviation of 150 bp.

With the similar pre-processing procedures on HIV 5 strains data, we used PEHaplo to assemble contigs from these data sets and compared the results with SAVAGE, which has better performance than MLEHaplo and IVA. The results are shown in Table 3.

The results reveal that both tools failed to assemble the rare strain with 5% or 1% abundance. However, PEHaplo was able to better assemble the dominant strain with one long contig. In addition, when the rare strain reached 10% (100x) of the total coverage, PEHaplo could partially assemble it, while SAVAGE only assembled the dominant one.

**Table 3.** Assembly results on simulated biased HXB2-NL43 MiSeq data set for SAVAGE and PEHaplo.

| HXB2<br>:NL43 | Tools | Contig<br>num | N50 | Genome<br>covered (%) | Unaligned<br>length (bp) | Mismatch<br>rate (%) | Indels<br>(%) |
| --- | --- | --- | --- | --- | --- | --- | --- |
| 900<br>:100 | SAVAGE | 7 | 2,500 | 46.76 | 581 | 0.022 | 0 |
|  | PEHaplo | 11 | 8,163 | 80.26 | 0 | 0.038 | 0 |
| 950<br>:50 | SAVAGE | 8 | 8,032 | 46.76 | 1,817 | 0 | 0 |
|  | PEHaplo | 1 | 9,470 | 46.76 | 0 | 0.033 | 0.01 |
| 990<br>:10 | SAVAGE | 13 | 2,130 | 46.75 | 1,590 | 0.022 | 0 |
|  | PEHaplo | 1 | 9,509 | 48.95 | 0 | 0 | 0 |

### 3.6 Benchmark on Influenza data set

In addition to HIV data, we also applied PEHaplo on a real Influenza H1N1 data set (SRR1766219) sequenced from the mix of a wild type (99%) and a mutant type (1%). This data set is sequenced with Illumina MiSeq sequencing technology, containing 646,879 pairs (2×250) of reads that cover the two strains to ∼23,000x. The mutant type carries two silent mutations in the M1 ORF (C354T and A645T, segment 7).

We first perform similar pre-processing on the Influenza data. With 851,988 filtered and error corrected reads, 27,888 reads were kept after removing duplicates and substrings. We still kept those reads that duplicate at least three times in the raw data set. After pre-processing, 11,940 reads were kept for strand adjustment and assembly.

It is worth noting that the H1N1 viruses have 8 segmented genomes. PEHaplo produced 10 contigs from the MiSeq Influenza data, with 8 contigs covering over 99% of the 8 segments of Influenza genome and 2 contigs unaligned. On the other hand, SAVAGE produced 220 contigs with a N50 value of 620 bp. The results are summarized and compared in Table 4. The comparison shows that PEHaplo works much better than SAVAGE on this Influenza quasispecies data as it successfully assembled all the 8 segments.

This data set contains a wild type and a rare mutant type (1%). However, neither method can recover the two mutations in the rare haplotype. In order to investigate this issue, we mapped all reads back to the region of the rare haplotype that contains the two mutations. The read mapping results clearly show that only several reads contain the same bases as the mutant type while all the other reads support the wild type. Thus, with such low number of mapped reads, existing information is not sufficient to 4 Discussion and conclusion distinguish true mutations from sequencing errors. Long read sequencing platforms might be a better choice for recovering the rare mutant type.

**Table 4.** Assembly results on Influenza MiSeq data set for SAVAGE and PEHaplo.

| Tools | Contig<br>num | N50 | Genomes<br>covered (%) | Unaligned<br>length (bp) | Mismatch<br>rate (%) | Indels<br>(%) |
| --- | --- | --- | --- | --- | --- | --- |
| SAVAGE | 220 | 620 | 96.3 | 38,303 | 0.818 | 0.046 |
| PEHaplo | 10 | 1,790 | 99.5 | 1,270 | 0.836 | 0.007 |

### 3.7 Computational time and memory usage

To evaluate the computational efficiency of our tool, we compare the running time and peak memory usage of the tested tools on the HIV 5-strain simulated data and also the real data. The results are shown in Table 5. PEHaplo runs significantly faster than SAVAGE and MLEHaplo. All the experiments were tested on a MSU HPCC CentOS 6.8 node with Two 2.4Ghz 14-core Intel Xeon E5-2680v4 CPUs and 128GB memory. We used 4 threads for IVA, 16 threads for SAVAGE and 1 thread for PEHaplo. The commands of running these tools on HIV simulated data can be found in Supplementary Materials Section 5.

**Table 5.** Running time and peak memory usage of assembly tools on HIV simulated and real data.

| Tools | Simulated data |  | Real data |  |
| --- | --- | --- | --- | --- |
|  | Time | Memory (GB) | Time | Memory (GB) |
| IVA | 17m | 2.6 | 17m | 2.6 |
| MLEHaplo | 10h | 6.4 | 4h | 2.4 |
| SAVAGE | 6h | 1.5 | 125h | 1.1 |
| PEHaplo | 9m | 2.9 | 23m | 1.3 |

## 4 Discussion and conclusion

For paired-end reads, one may consider to combine read pairs into a longer sequence before conducting assembly. We actually applied existing read joining tools for this purpose. However, joining reads is not a trivial problem as the overlapping part of the read pairs may not always be identical. Thus, existing methods of joining two ends may introduce errors. In addition, merging paired-end reads will discard the paired-end information for guiding the path finding process. As a result, the experimental results using PEAR (Zhang *et al*., 2014) and other end merging tools show inferior performance. Thus we did not include that step in our pipeline.

The third-generation sequencing platforms such as PacBio can produce very long reads, which can cover the whole length of viral genomes. However, the high sequencing error rate (about 10%) and the lower throughput than Illumina still hamper their wide application for metagenomic sequencing. The advantages and limitations of applying current long reads technologies for viral haplotypes reconstruction are discussed in BAsE-Seq (Hong *et al*., 2014). With the increased read quality, long read sequencing technologies will greatly simplify the assembly methods for metagenomic data (Di Giallonardo *et al*., 2014). However, at this moment, viral haplotype reconstruction using short reads is still needed.

Our method can be extended to metagenomic data if the member species’ genomes have common regions with length smaller than fragment size. However, our analysis has shown that many genes in metagenomic data can have LCS sizes much greater than typical fragment size. For those metagenomic data, large insert sizes should be chosen for the sequencing protocol.

In conclusion, we presented a de novo viral haplotype reconstruction tool for viral quasispecies. We applied it to both simulated and real quasispecies data and achieved better results than several benchmarked tools.

## Acknowledgements

We would like to acknowledge the help from Nan Du on producing some figures in this manuscript.

